# Gene therapy-mediated overexpression of wild-type MFN2 improves Charcot-Marie-Tooth disease type 2A

**DOI:** 10.1101/2025.10.15.682364

**Authors:** Marine Tessier, Zeinab Hamze, Nathalie Bonello-Palot, Nathalie Roeckel-Trévisiol, Shahram Attarian, Marc Bartoli, Valérie Delague, Bernard L. Schneider, Nathalie Bernard-Marissal

## Abstract

Charcot-Marie-Tooth disease type 2A (CMT2A) is the most common axonal CMT and is associated with an early onset and severe motor-dominant phenotype. CMT2A is mainly caused by dominant mutations in the *MFN2* gene, encoding Mitofusin-2, a GTPase located in the outer membrane of the mitochondria and endoplasmic reticulum (ER). Mutations in *MFN2* are known to affect mitochondrial dynamics. We previously demonstrated that the mutated MFN2^Arg94Gln^ further disrupts contacts between the ER and the mitochondria, leading to progressive axonal degeneration. There is no effective therapeutic approach to slow or reverse the progression of CMT2A, and treatments currently under development primarily focus on restoring mitochondrial function.

Here, we provide proof-of-concept that neuronal overexpression of wild-type MFN2 (MFN2^WT^) provides therapeutic benefit in transgenic CMT2A mice carrying the mutated MFN2^Arg94Gln^. Intrathecal delivery of an AAV9 vector expressing MFN2^WT^ effectively targets motor and sensory neurons, restoring ER-mitochondria contacts and mitochondrial morphology, thereby preserving both neuromuscular junction integrity and motor function. Strikingly, therapeutic efficacy is also achieved following vector injection after the onset of symptoms, rescuing the molecular hallmarks of CMT2A pathology and reversing locomotor. Notably, AAV administration was well tolerated, with no evidence neither of hepatotoxicity nor dorsal root ganglion inflammation. These results establish that boosting MFN2’s levels using gene therapy is a promising therapeutic avenue for CMT2A.

## Introduction

Charcot-Marie-Tooth (CMT) disease is one of the most common inherited neuromuscular diseases affecting the peripheral nervous system. With a prevalence of 1 in 2,500 to 10,000^1^, it is estimated that ca. 300,000 people suffer from CMT in Europe (https://ecmtf.org/about-cmt/) and ca. 2.6 million worldwide (https://www.hnf-cure.org/cmt/what-is-cmt/). Disease severity varies considerably between subtypes. Symptoms typically appear within the first two decades of life^2^. Most of the patients display gait disorders due to distal amyotrophy of the lower limbs, often accompanied by sensory alterations. In the most severe cases, patients become wheelchair-dependent^1^. More than 100 subtypes of CMT exist according to the causal genes and mode of inheritance and cell-type affected (Schwann cells or in motor and sensory neurons)^1^. This heterogeneity complicates therapeutic development^3^ with most therapies currently focusing on CMT1A, the most prevalent form accounting for 70% of all CMTs^1^. In consequence, CMT disease remains incurable and only symptomatic and supportive cares can be provided to patients. Hence, the economic impact of CMT remains significant with an estimate cost of approximately $22,000/year per patient in Western European countries^4^.

CMT2A is the most common form of axonal CMT, accounting for 5-10% of all CMT^5^. It is often associated with an early-onset severe motor-dominant neuropathy, which may be accompanied by optic nerve atrophy, scoliosis, hearing loss, vocal cord paralysis and upper motor neuron involvement^5^. CMT2A is mainly caused by autosomal dominant heterozygous mutations (over 100) in the *MFN2* gene which encodes the Mitofusin-2 protein (MFN2)^5–7^. MFN2, like its counterpart MFN1, is a GTPase anchored to the outer mitochondrial membrane, where it regulates mitochondrial fusion through the formation of homo- (with MFN2) or heterodimers (with MFN1). MFN2 is also present on the endoplasmic reticulum (ER) surface, enabling close contacts between the ER and mitochondrial membranes^8,9^. These contacts regulate various processes including lipid metabolism, calcium homeostasis, mitochondrial dynamics, and autophagy/mitophagy, which are essential for neuronal and axonal function^10^. We previously highlighted that the *MFN2*^*Arg94Gln*^ mutation disrupts contacts between ER and mitochondria leading to impaired mitochondrial dynamics, ER stress and axonal degeneration largely contributing to CMT2A pathology^11^. Since then, other groups have reported defective contacts between these organelles in different cell models, as well as related mutations in the *MFN2* gene^12–14^.Therefore, we consider that improving the interplay between the ER and the mitochondria could be a valuable therapeutic approach to counteract CMT2A pathology. However, current therapeutic approaches for CMT2A mainly focus on restoring mitochondrial function, either by using synthetic compounds that promote mitochondrial fusion, or overexpressing MFN1, the homolog of MFN2^15–18^. Another option is the development of HDAC6 inhibitors to counteract loss of acetylated tubulin^19,20^. Although appealing, those strategies may have limited effectiveness and potential adverse effects as they fail to address the dual dysfunction of mitochondria and ER, and do not specifically target the most affected cells, namely motor and sensory neurons.

Conversely, therapy targeted to the affected cell type is possible thanks to gene therapy by selecting appropriate vector, serotype, expression cassette and route of administration^21,22^. In recent years, there has been increased development of gene therapy in the field of CMT^23^, with the launch of the first clinical trials for giant axonal neuropathy^24^ (NCT01503125) and IGHMBP2-related neuropathies (NCT05152823). For CMT2A, a proof of concept has been reported combining RNA interference with gene replacement *in vitro*. This strategy would allow to target multiple mutated allele linked to CMT2A, but may require the use of dual AAV *in vivo* that complicates therapeutic development^25^. Here, we present the first *in vivo* gene therapy approach for CMT2A. We demonstrate that the overexpression of the wild-type (WT) form of MFN2, targeted specifically to motor and sensory neurons using AAV9 under the control of the human synapsin1 promoter, can counteract CMT2A pathology, regardless of whether the treatment is administered at the pre- or post-symptomatic stage. Additionally, we show that overexpression of MFN2^WT^restores ER-mitochondria contacts, indicating that therapeutic benefits can be achieved by restoring the interplay between these organelles.

## Materials and methods

### Constructs and production of AAV vectors

pAAV-hsyn-MFN2^WT^ was previously generated in^11^. The various constructs were packaged into AAV6 and AAV9 particles at the EPFL Bertarelli platform for gene therapy (Geneva, Switzerland). For each vector the viral genome concentration was determined by dPCR (QIAcuity apparatus, QIAGEN), using an amplicon located in the WPRE sequence. For AAV6, the infectivity titer of each virus (expressed as Transduced Units/mL) was determined following infection of HEK293T cells by real-time PCR using primers for WPRE element (forward: 5’ CCG TTG TCA GGC AAC GTG 3’, reverse: 5’AGC TGA CAG GTG GTG GCA AT 3’, probe: FAM-TGC TGA CGC AAC CCC CAC TGG T-BHQ1) and human albumin (forward: 5’ TGA AAC ATA CGT TCC CAA AGA GTT T 3’, reverse: 5’ CTC TCC TTC TCA GAA AGT GTG CAT AT 3’, probe: FAM-TGC TGA AAC ATT CAC CTT CCA TGC AGA-TAMRA).

### Animals

*In vivo* work was performed using the mouse strain B6;D2-Tg (Eno2-MFN2*R94Q) L51Ugfm/J previously described^5^ (alternative name MitoCharc1, purchased from Jackson Laboratories, stock No: 012812). Animals were maintained as heterozygous by crossing B6;D2-Tg (Eno2-MFN2*R94Q)/-males with B6;D2F1 females (Charles river). Offspring was genotyped using a standard PCR reaction with the following primers recommended by Jackson Laboratories: 10453: 5’ ATG CAT CCC CAC TTA AGC AC 3 3’, 10454: 5’ CCA GAG GGC AGA ACT TTG TC 3’; oIMR8744: 5’ CAA ATG TTG CTT GTC TGG TG 3’, oIMR8745: 5’ GTC AGT CGA GTG CAC AGT TT 3’. Animals were housed in an animal facility with 12 h light /12 h dark environment, and ad libitum access to water and normal diet. All animal experiments complied with the French ethical committee, which approved all experimental procedures (No: APAFIS#37715), and the ARRIVE guidelines and were carried out under French and EU regulations (2010/63/EU and national decree n°213-118), following recommendations of the local ethics committee.

### Intrathecal Injection

Intrathecal injections were performed to deliver viral vectors into the cerebrospinal fluid at lumbar level of adult mice (2- or 4.5-month-old mice). Mice were anesthetized with isoflurane (4% for induction, 1.5–2% for maintenance) and injected in subcutaneous with buprenorphine (0.1mg/kg) to prevent any pain and positioned prone on a heating pad. The lumbar area was shaved. Using a 30G insulin syringe, a 10-15⍰μL volume of viral suspension was injected rapidly into the L5–L6 intervertebral space. The total dose of AAV9-hSyn1-GFP or MFN2 vectors injected was 5^E^11 viral genomes (VG). We had previously tested a lower dose of 2^E^11 VG, but the proportion of motor neurons transduced was low (less than 20%) compared to the 5^E^11 VG dose. Successful intrathecal entry was indicated by a reflexive tail flick upon needle insertion. The needle was withdrawn immediately after injection. Mice were monitored until full recovery from anesthesia and then returned to their home cages. The animals were carefully examined every day during the week following the injection.

### Tissue Preparation

Mice were sacrificed by perfusion intracardially with cold phosphate buffer saline (PBS 1X) prior to a 4% paraformaldehyde solution (diluted from a 32% PFA solution, Thermofischer, 047377.9L). Tissue samples (Lumbar spinal cord and Tibial Nerve), were fixed by immersion in 4% PFA in 0.⍰1M phosphate-buffered saline (PBS) for 24 hours at 4⍰°C, then cryoprotected in 20% (w/v) sucrose solution until embedding. Samples were embedded in Optimal Cutting Temperature compound (OCT, TFM™) and snap-frozen at −80⍰°C. Serial cryosections (14μm thick for nerves, DRG and lumbrical and 20 µm for spinal cord) were cut using a cryostat (Leica CM1860) and mounted on Superfrost Plus slides (Thermo Fisher Scientific). Slides were stored at −20⍰°C until immunostaining.

### Immunofluorescence Staining

Spinal cord, nerves, and DRG sections were permeabilized and blocked for 1 h at room temperature in PBS containing 0.3% Triton X-100 and 5% fetal bovine serum (FBS). Sections were then incubated overnight at 4 °C with primary antibodies diluted in PBS containing 0.3% Triton X-100 and 5% FBS: rabbit anti-stathmin-2 (NBP1-49461, Novusbio, 1:500), chicken anti-NFM (822701, Biolegend, 1:500), mouse anti-NeuN (AB104224-1001, Abcam, 1:1000), mouse and rabbit anti-GFP (mouse: 11814460001, Roche; rabbit: A11122, Invitrogen, 1:500), rabbit anti-Iba1 (019-19741, Wako, 1:500), and rabbit anti-ATF3 (33593T, Cell Signaling, 1:500). Lumbrical sections were processed similarly, using 0.5% Triton X-100 and 5% FBS for permeabilization and blocking. After washing (3 × 5 min in PBS), sections were incubated for 1 h at room temperature with fluorophore-conjugated secondary antibodies (Alexa Fluor®, Abcam, 1:500) protected from light. Following additional washes, slides were mounted with DAPI-containing antifade medium (ProLong™ Diamond Antifade Mountant) and stored at 4 °C until imaging. Imaging was performed using an ApoTome structured illumination system (Zeiss) in Z-stack mode to enhance contrast and optical sectioning at 10X magnification. For stathmin-2 labeling, fluorescence intensity was quantified using Corrected Total Cell Fluorescence (CTCF), calculated as: CTCF=Integrated Density − (Area of selected cell × Mean fluorescence of background). CTCF was used as it corrects for background signal and provides a more accurate measure of fluorescence intensity, allowing reliable comparisons across samples.

### Electron microscopy

Mice were perfused intracardially with cold phosphate buffer saline (PBS 1X) prior to a 4% paraformaldehyde solution (diluted from a 32% PFA solution, Thermofischer, 047377.9L). Lumbar spinal cords and tibial nerves were dissected out and transferred in a fixative solution (PFA 2% + 2.5% Glutaraldehyde in 0.1M cacodylate buffer, Thermofischer) overnight for the SCL and 6h for the nerves at 4 °C. Mice were perfused intracardially with cold phosphate buffer saline (PBS 1X) prior to a 4% paraformaldehyde solution (diluted from a 32% PFA solution, Thermofischer, 047377.9L). Lumbar spinal cords and tibial nerves were dissected out and transferred in a fixative solution (PFA 2% + 2.5% Glutaraldehyde in 0.1M cacodylate buffer, Thermofischer) overnight for the SCL and 6h for the nerves at 4 °C. The tissues were then washed three times in 0.1M cacodylate buffer and post-fixed in buffered 1% OsO4 for one hour at 4°C. After washes in distilled water, the samples were contrasted in aqueous 1% uranyl acetate. Samples were then dehydrated in graded series of ethanol baths (30 minutes each) at 4°C and infiltrated with epon resin in ethanol (1:3, 2:2, 3:1) for 2 hours for each, and finally in pure resin overnight. The next day the spinal cord and nerves were embedded in fresh pure epon resin and cured for 48h at 60°C. 1000 nm semi-thin and 80 nm ultra-thin sections were performed on a Leica UCT Ultramicrotome (Leica, Austria). Semi-thin sections were stained with toluidine blue and ultrathin sections were deposited on formvar-coated slot grids. The grids were contrasted using lead citrate (5 minutes) and observed using an FEI Tecnai G2 at 200 KeV. The acquisition was performed on a Veleta camera (Olympus, Japan).

The number of mitochondria connected to the ER, was evaluated in the lumbar spinal cord of 5- or 12-month-old WT and CMT2A Tg mice treated with either PBS or the AAV vectors. Mitochondria was considered as connected to the ER if the distance between the two organelles was below 40nm. The number of attached mitochondria was expressed in percentage relative to the total number of mitochondria per field as described in ^11^.

### Viral genome copies number quantification

Fresh tissues (spinal cord, sciatic and tibial nerves, DRG and liver) were collected 3 months post-injection of CMT2A mice treated with AAV-hsyn-GFP or MFN2^WT^. Samples were lysed using a tissue lyser (TissueLyser II, Qiagen) and total DNA was extracted using the Maxwell® CSC benchtop automated instrument (Promega) with a viral DNA extraction kit (Promega, AS1330). The dPCR assays were performed on a QIAcuity instrument (QIAGEN), using the QIAcuity× PCR kit in 8.5k 96-well nanoplates. For dPCR measurements of vDNA abundance, we used an amplicon present in the WPRE vector sequence. The total amount of gDNA input was determined with an amplicon in the mouse albumin gene. We considered that two albumin copies represented one diploid mouse genome, which allowed to determine an estimated number of episomal vector copies per cell.

### Behavioral Studies

Locomotion was evaluated by the rotarod test. Motor coordination and balance were evaluated using an accelerating (4-40 rpm, 5 minutes) rotarod apparatus (TSE systems). The protocol consisted of three phases over 3 days as described in ^11^:

Before each session, the mice were placed in the experimental room 30 minutes before the start of the session in order to get used to their new environment.

#### Day 1 (Habituation)

Mice were placed individually on the stationary rotarod to freely explore the apparatus for 5 minutes to minimize stress and familiarize them with the device.

#### Day 2 (Training)

Each mouse underwent three timed trials on the rotarod, with the rod accelerating from 4 rpm to 40 rpm over 5 minutes. The latency to fall (time in seconds before the mouse fell off or completed a full rotation while hanging on) was recorded automatically. A rest period of at least 15 minutes was allowed between trials.

#### Day 3 (Evaluation)

After one day of rest, mice were tested again with three timed trials under the same acceleration conditions as during training. The maximal latency to fall across the three trials was calculated for each mouse and used for subsequent analysis.

In addition, the mice were handled and weighed weekly.

## Statistical analysis

*In vivo* experiments were designed and conducted in accordance with the recommendations of the ARRIVE guidelines. Animals cohorts were matched according to gender, genotype and age. Mice were randomly assigned to experimental groups and all behavioral experiments as well as processing and analyzes of the tissues were performed in blind. Size of the cohort (7 - 10 animals per group) for behavioral studies was estimated based on previous experiments using the CMT2A Tg model and by taking into account an alpha risk of less than or equal to 0.05, a test power of 90% and a moderate effect of 30% (http://www.lasec.cuhk.edu.hk/sample-size-calculation.html). We anticipated to increase the size of each group by 20 % in order to anticipate the potential loss of animals during the experiment. Indeed, in the rotarod test, some mice were excluded from the analysis as they did not move on the rod during the first 3 sessions or died during the study for reasons independent from the treatment (for early treatment experiment: 1 WT+PBS, 2 CMT2A+GFP and 3 CMT2A+MFN2^WT^ were excluded; for late treatment: 1 WT+PBS and 1 CMT2A+MFN2^WT^ were excluded). For histology, we analyzed 3 to 6 mice per condition.

In most of the conditions, ordinary one-way ANOVA test with multiple comparison and Tukey’s post hoc test were used to compare three or more groups. When variances were unequal, the Brown– Forsythe ANOVA was used. All mean values are given with the standard error of the mean (SEM). When the number of values per condition was superior to 30, a normality test (Shapiro-Wilk) was applied. When the normality assumption was violated, the Kruskal–Wallis multiple comparison with Dunn’s post hoc test was applied. In this case, values are expressed as median with the min and max. For all experiments, the detailed number of n and the statistical test applied are defined in the figure captions. Statistical analyses were carried out using Prism Version 8.3.0 (GraphPad software, Inc., La Jolla, CA, USA) and are represented as follows: *P < 0.05, **P < 0.01 and ***P < 0.001. The applied statistical tests as well as the number of replicates are indicated in the figure legends.

## Data availability

The authors confirm that the data supporting the findings of this study are available within the article and/or its Supplementary material. These data are available from the corresponding author, upon request.

## Results

### Intrathecal injections of AAV9-hSyn1-GFP and MFN2 ^WT^ target both motor neurons in the lumbar spinal cord and sensory neurons in the DRGs

Despite the predominance of motor symptoms, CMT2A is known to affect both sensory and motor neurons (MNs)^5^. To specifically counteract the effect of the MFN2^Arg94Gln^ in affected neurons, we used AAV9 combined with the hsyn1 promoter to express either GFP or MFN2^WT^. AAV9-hsyn1-GFP, MFN2^WT^ or PBS were delivered to 2-month-old CMT2A via intrathecal lumbar injections (**Figure 1A-B**). Wildtype (WT) mice were injected only with PBS as control. Expression of the transgenes (i.e. GFP or myc-MFN2) was analyzed 3 months later at the onset of locomotor impairments in CMT2A Tg mice. We observed a high level of transduction of lumbar MNs in both AAV9-hsyn1-GFP and MFN2^WT^ groups (GFP^+^ MNs: 59.3±6.2%; MFN2^+^ (myc+) MNs: 57.7±4.7%). Transduction efficiency was lower in sensory neurons (GFP^+^ MNs: 23.2±4.9%; MFN2^+^ (myc+) MNs: 21.06±3.5%) (**Figure 1C-E**). We also observed the presence of a transgene-expressing cells in the cervical and thoracic spinal cord, as well as in the brain, mainly in areas close to the ventricles, such as the hippocampus (**Supplementary Figure 1. A-B**).

**Figure 1.**
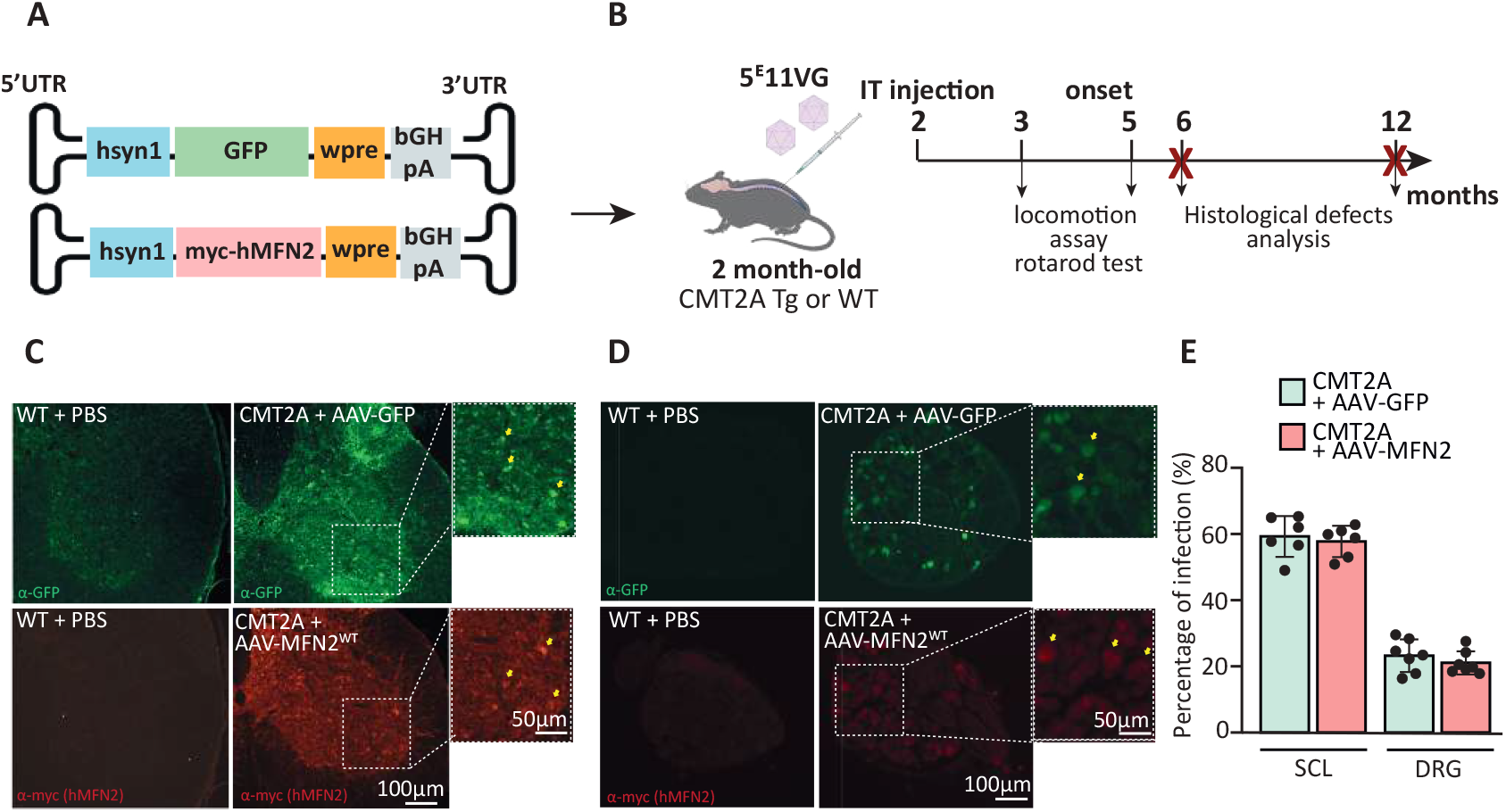
Spinal cord and DRG targeting after intrathecal injections of AAV9-hsyn vectors. (A) Experimental AAV vector design: hsyn1 promoter drives expression of GFP or myc-hMFN2^WT^ (NM_014874.4, myc tag was added in N-terminal). (B) Experimental design: 2-month-old CMT2A transgenic (Tg) or wild-type (WT) mice received intrathecal injections at the lumbar level of 5^E^11 vg AAV-hsyn-GFP or AAV-hsyn-MFN2^WT^ or PBS. Motor dysfunction was assessed at 3–5 months using rotarod test, and histological analyses were performed at 3- and 10-months post-injection, corresponding to 5- and 12-month-old animals respectively. (C) Representative immunofluorescent staining of lumbar spinal cord sections showing GFP (green, top) or myc-hMFN2 (red, bottom) expression following AAV administration (right panels) compared with WT + PBS controls (left panel). Higher magnifications of the ventral horn are depicted where yellow arrows indicate transduced motor neurons. (D) Representative images of dorsal root ganglia (DRG) sections showing GFP (green, top) or myc-hMFN2 (red, bottom) expression following AAV injection (right panels) compared with WT + PBS controls (left panels. Insets highlight transduced neurons (yellow arrows). (E) Quantification of transduction efficiency in lumbar spinal cord (SCL) and DRG, showing comparable infection rates between AAV-GFP and AAV-hMFN2 (N=6 animals per condition). Data are presented as mean ± SEM. (A and B) has been created with Biorender.

### Overexpression of MFN2 ^WT^ at pre-symptomatic stage prevents locomotor defects and axonal degeneration in CMT2A Tg mice

We previously showed that the MFN2^Arg94Gln^ mutation results in impaired mitochondrial dynamics and ER-mitochondria contacts, similar to the consequence of MFN2 loss ^8,9,11^. We therefore hypothesized that enhancing MFN2^WT^ levels in affected neurons could promote functional MFN2^WT^ homodimerization, counteracting the effects of MFN2^Arg94Gln^, and consequently CMT2A pathology. In a previous proof-of-concept study, we observed that MFN2^WT^ overexpression prevented axonal degeneration in primary mouse CMT2A MNs *in vitro*^11^. To determine whether this strategy could be beneficial *in vivo*, we delivered our therapeutic AAV vector (i.e. AAV9-hsyn1-MFN2^WT^) or controls vectors (AAV9-hsyn1-GFP) or PBS in 2-month-old pre-symptomatic CMT2A mice. WT mice were injected with PBS at the same age. Using the rotarod test, we monitored the locomotion of the mice before (3 months) and after (5 months) the onset of motor symptoms. We did not observe any difference between the four groups at 3 months, demonstrating that the treatment was well tolerated (WT + PBS: 134.8±18.11 sec, CMT2A+PBS: 143.1±19.2 sec, CMT2A+ AAV9-hsyn1-GFP: 125.3±18.9 sec and CMT2A+ AAV9-hsyn1-MFN2^WT^: 160.1±14.5 sec) (**Figure 2A)**. As expected, at 5 months of age, CMT2A mice injected with PBS and AAV9-hsyn1-GFP showed a significant decrease in the latency on the rod, compared with WT mice treated with PBS (WT + PBS: 125.5±7.6 sec, CMT2A+PBS: 86.2±7.9 sec, CMT2A+ AAV9-hsyn1-GFP: 93.4±9.2 sec) (**Figure 2B**). Interestingly, CMT2A mice injected with AAV9-hsyn1-MFN2^WT^ displayed a similar latency on the rod to WT mice (CMT2A+AAV9-hsyn1-MFN2^WT^: 127.8±10.4 sec) (**Figure 2B)**. The motor function of the mice injected with AAV9-hsyn1-MFN2^WT^ was significantly improved compared to CMT2A+PBS mice, and there was a strong trend towards improvement compared to CMT2A+AAV-hsyn-GFP mice (p:0.06). We then determined whether this protective effect of AAV9-hsyn1-MFN2^WT^ on locomotion was associated with prevention of axonal abnormalities. In particular, we analyzed the expression of Stathmin-2, a microtubule-binding protein, reported to be upregulated in the nerves of a CMT2A rat model^26^. We confirmed the upregulation of stathmin-2 levels in the tibial nerves of the CMT2A mice injected with either PBS or AAV9-hsyn1-GFP, compared to WT mice (WT + PBS: 3.51±0,6^E^6 a.u, CMT2A+PBS: 2.03±0,5^E^7 a.u, CMT2A+AAV9-hsyn1-GFP: 1.5±0,3^E^7 a.u). Importantly, this upregulation was prevented by overexpressing MFN2^WT^ (CMT2A+AAV9-hsyn1-MFN2^WT^: 4.25 ±1.2^E^6) (**Figure 2C-D**). Finally, we analyzed muscle denervation of the lumbrical muscles in the hind paws. We observed a significant increase in the proportion of partially denervated NMJs (i.e. intermediate) in CMT2A mice treated with AAV9-hsyn1-GFP or PBS compared to WT mice. This increase was prevented by treatment with AAV9-hsyn1-MFN2^WT^ (intermediate NMJs: WT + PBS: 15.9±1.7%, CMT2A + PBS: 33.7±6.7%, CMT2A +AAV9-hsyn1-GFP: 32.8±1.6% and CMT2A +AAV9-hsyn1-MFN2^WT^: 16.9±3.3%) (**Figure 2E-F)**. In conclusion the overexpression of MFN2^WT^ prevents the development of locomotor defects and axonal degeneration associated with CMT2A pathology.

**Figure 2.**
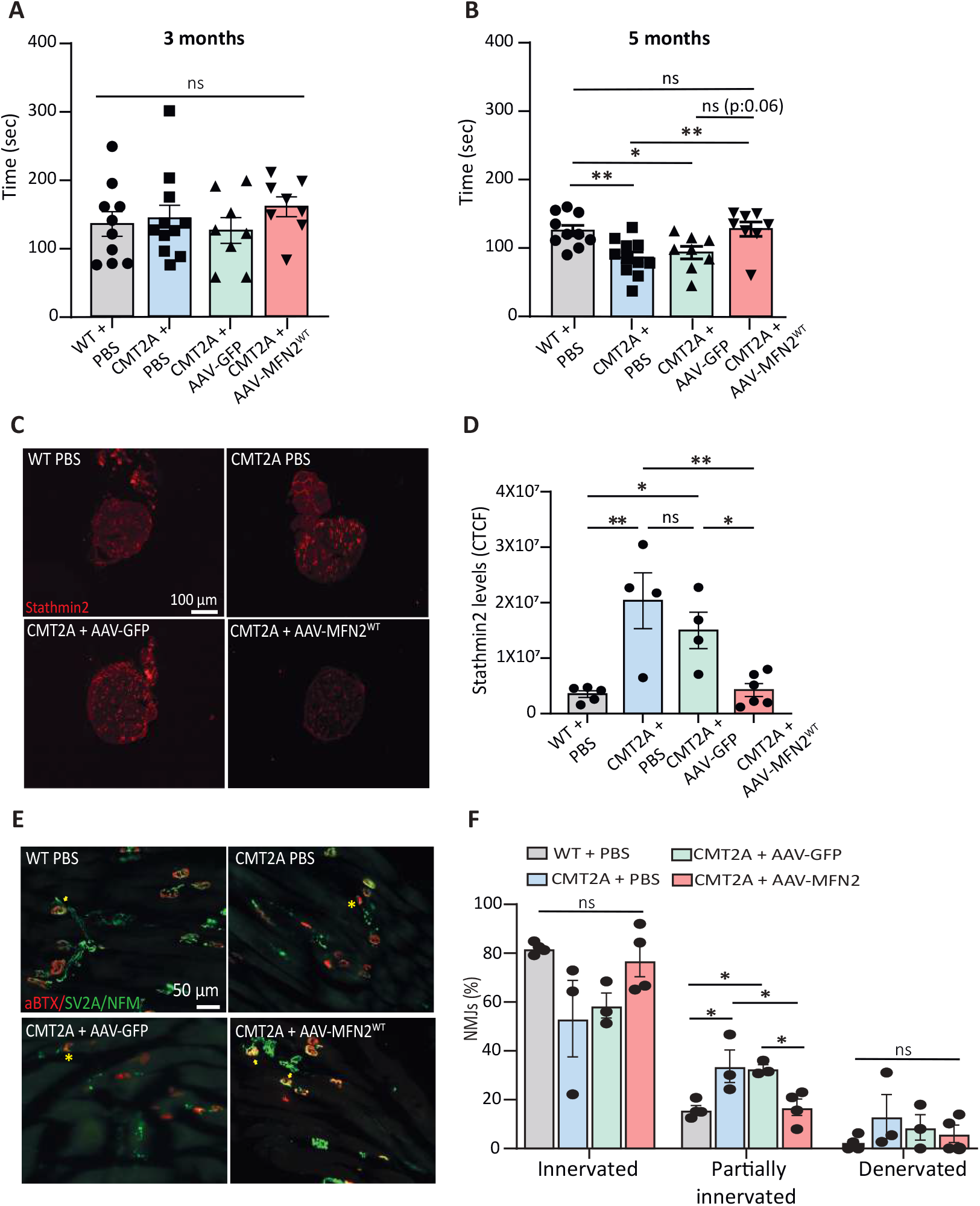
Pre-symptomatic overexpression of MFN2 ^WT^ prevents locomotion defects and axonal degeneration in CMT2A Tg mice. (A) Evaluation of the performance on the rotarod at 3 months old. CMT2A Tg mice do not show impaired motor disabilities at this stage. (B) Evaluation of the performance on the rotarod at 5 months old. For both 3- and 5-month-old time points, experimental groups were distributed as followed: WT+PBS: n=10; CMT2A+PBS: n=11; CMT2A+AAV-GFP: n=8 and CMT2A+AAV-MFN2^WT^: n=8. Data are shown as mean ± SEM. Statistical analysis: one-way ANOVA multiple comparisons with Tukey’s post hoc test. Motor deficits observed in CMT2A Tg mice are significantly prevented by AAV-hMFN2 treatment. (C) Representative immunofluorescence staining of transverse sections of tibial nerves of 6-month-old treated mice using anti-stathmin2 (red) (D) Quantification of stathmin2 intensities in transverse sections of tibial nerves using corrected total cell fluorescence (CTCF) analysis. Stathmin2 levels are reduced in the nerves of CMT2A Tg mice and restored following AAV-hMFN2 administration (WT+PBS: n=5; CMT2A+PBS: n=4; CMT2A+AAV-GFP: n=4 and CMT2A+AAV-MFN2^WT^: n=6). Data are shown as mean ± SEM. Statistical analysis: one-way ANOVA multiple comparisons with Tukey’s post hoc test.(E) Representative images of staining of neuromuscular junctions in lumbrical muscles of 12-month-old treated mice labeled with α-bungarotoxin (α-bungarotoxine, BTX, red) and anti-SV2A and NFM (green) (F) Quantification of NMJ innervation status (WT+PBS: n=4; CMT2A+PBS: n=3; CMT2A+AAV-GFP: n=3 and CMT2A+AAV-MFN2^WT^: n=4). Note the presence of denervated NMJs positive for αBTX only indicated by asterisk; innervated NMJs are positive for both αBTX anti-SV2A and NFM and indicated by yellow arrows. Data are shown as mean ± SEM. Statistical analysis: one-way ANOVA multiple comparisons with Tukey’s post hoc test. For all conditions, *p* < 0.05 was considered significant.

### Overexpression of MFN2 ^WT^ restores ER-mitochondria contacts in CMT2A Tg mice

We and others have previously demonstrated that the disconnection between the ER and mitochondria is key for the development of CMT2A pathology^11,14^. We therefore analyzed in MNs of the lumbar spinal cord, using electron microscopy, whether the therapeutic effect of MFN2^WT^ overexpression was indeed linked to a rescue of ER and mitochondria interactions in MNs of WT and CMT2A mice. We observed that the overexpression of MFN2^WT^ was sufficient to restore the contacts that were reduced in CMT2A mice injected with PBS or the AAV9-hsyn1-GFP (WT+PBS: 85.5±10%, CMT2A+PBS: 25.8±5.6%, CMT2A+AAV9-hsyn1-GFP: 40.5±4.1% and CMT2A+AAV9-hsyn1-MFN2^WT^: 108.5±19.8%) **(Figure 3A-B)**. Since MFN2 is also known to regulate mitochondrial function, we analyzed in parallel the length of mitochondria in those MNs. Mitochondrial length was significantly reduced in the MNs of control CMT2A mice compared to the WT. However, the value of mitochondrial length was comparable to WT conditions in CMT2A mice treated with AAV9-hsyn1-MFN2^WT^ (WT+PBS: 0.69±0.02 µm, CMT2A+PBS: 0.53±0.02 µm, CMT2A+AAV9-hsyn1-GFP: 0.55±0.008 µm and CMT2A+AAV9-hsyn1-MFN2^WT^: 0.67±0.005 µm) **(Figure3A-C)**. Altogether, pre-symptomatic overexpression of preserves contacts between the ER and mitochondria as well as proper mitochondrial fusion.

**Figure 3.**
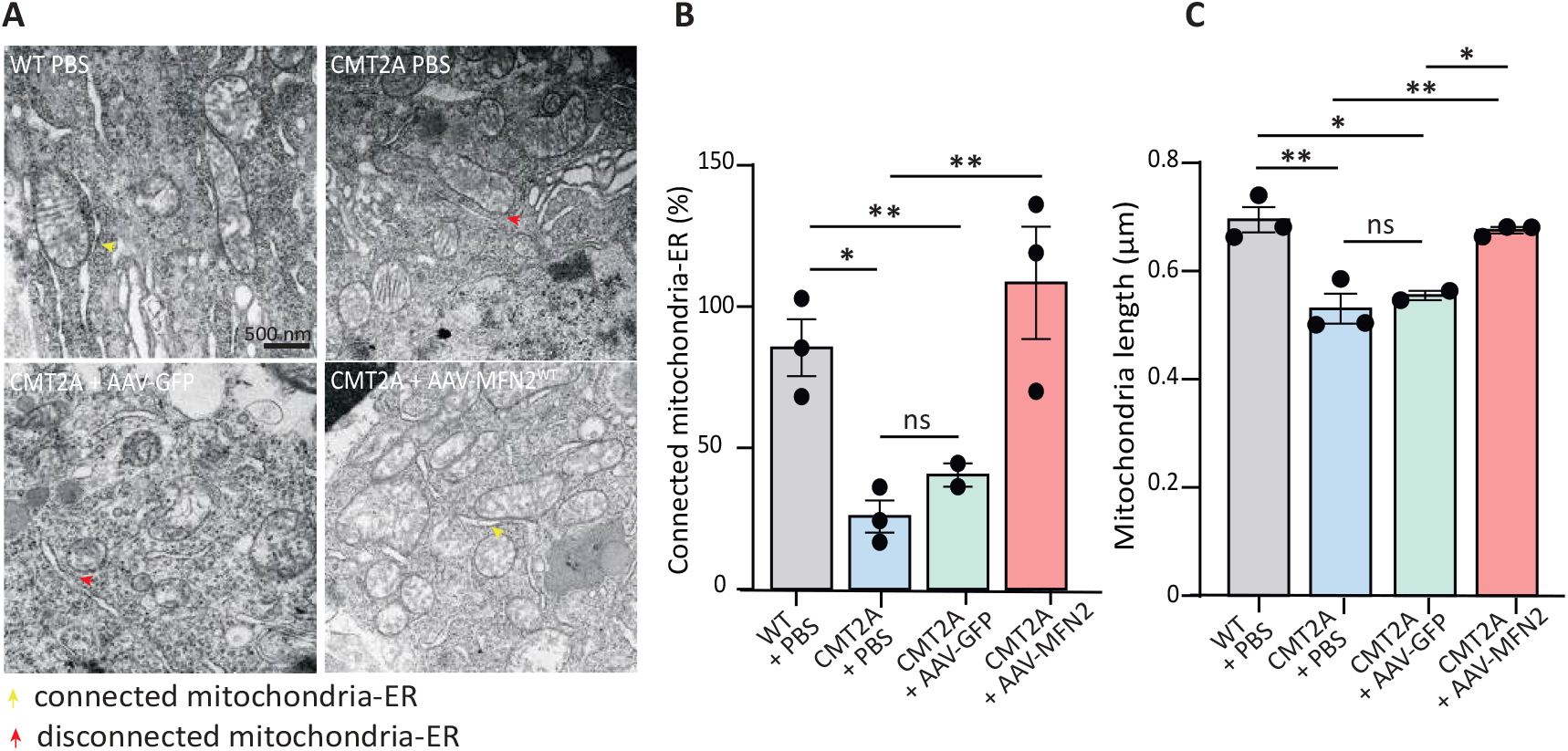
Early overexpression of MFN2 ^WT^ restores ER-mitochondria contacts and mitochondrial length. (A) Representative transmission electron microscopy (TEM) images illustrating the contacts between ER and the mitochondria in the soma of motor neurons of WT and CMT2A mice treated with PBS, AAV-GFP, or AAV-hMFN2^WT^. Yellow arrowheads indicate mitochondria in close apposition to the endoplasmic reticulum (ER), while red arrowheads highlight disconnected mitochondria–ER contacts. (B) Quantification of the percentage of mitochondria with ER contact in motor neuron soma in the lumbar spinal cord of 12-month-old mice. AAV-hMFN2 treatment significantly restores ER– mitochondria interactions in CMT2A mice. (C) Quantification of mitochondrial length in spinal cord motor neurons. For B and C, quantification was done on n=3 animals per group. Data are shown as mean ± SEM. Statistical analyses were performed using one-way ANOVA multiple comparisons with Tukey’s post hoc test; *p* < 0.05 was considered significant.

### Overexpression of MFN2 ^WT^ after symptom onset counteracts CMT2A pathology

Given that MFN2 overexpression has a preventive effect on the onset of CMT2A pathology, we sought to evaluate its protective effect once symptoms had appeared. To this end, we injected CMT2A mice with either the AAV9-hsyn1-GFP and AAV9-hsyn1-MFN2^WT^ at 4.5 months (18 weeks of age), a time point corresponding to the onset of the symptoms **(Figure 4A)**. We monitored locomotion before the injections (18 weeks of age), and 6 weeks post-injection (24 weeks of age, ∼6 months). At 4.5 months, CMT2A mice injected with AAV9-hsyn1-MFN2^WT^ initially displayed a significant alteration of their performance on the rod compared to WT mice (WT + PBS: 67.3±5.19 sec; CMT2A + AAV9-hsyn1-GFP: 52.4±4.1 sec; CMT2A + AAV9-hsyn1-MFN2^WT^: 44.9±4.2 sec) **(Figure 4B)**. Remarkably, the locomotor performance of the CMT2A mice treated with the AAV9-hsyn1-MFN2^WT^ was significantly and fully restored within 6 weeks compared to CMT2A mice injected with AAV9-hsyn1-GFP (CMT2A + AAV9-hsyn1-GFP: 52.3±7.4 sec; CMT2A + AAV9-hsyn1-MFN2^WT^: 86.9±5.3 sec) **(Figure 4B)**. As with pre-symptomatic treatment, this therapeutic effect was confirmed by restoring stathmin-2 expression levels in the tibial nerves of CMT2A mice overexpressing MFN2^WT^ compared with those overexpressing GFP (WT + PBS: 1.81±0.16^E^6 a.u; CMT2A + AAV9-hsyn1-GFP: 4.84±0.27^6^5 a.u; CMT2A + AAV9-hsyn1-MFN2^WT^: 1.63 ±0.25^E^6 a.u) **(Figure 4C-D)**. Finally, EM analysis of the spinal cord and nerves tissues confirmed that AAV-hsyn-MFN2 rescued the number of contacts between the ER and mitochondria in MNs of the spinal cord (WT + PBS: 130.7±5.42 %; CMT2A + AAV9-hsyn1-GFP: 71.7±10.28 %; CMT2A + AAV9-hsyn1-MFN2^WT^: 117.4±9.6 %) (**Figure 4E-G)** as well as mitochondrial length in tibial nerves (WT + PBS: 0.80±0.047 µm; CMT2A + AAV9-hsyn1-GFP: 0.53±0.068 µm; CMT2A + AAV9-hsyn1-MFN2^WT^: 0.79±0.057 µm) **(Figure 4H-I)**.

**Figure 4.**
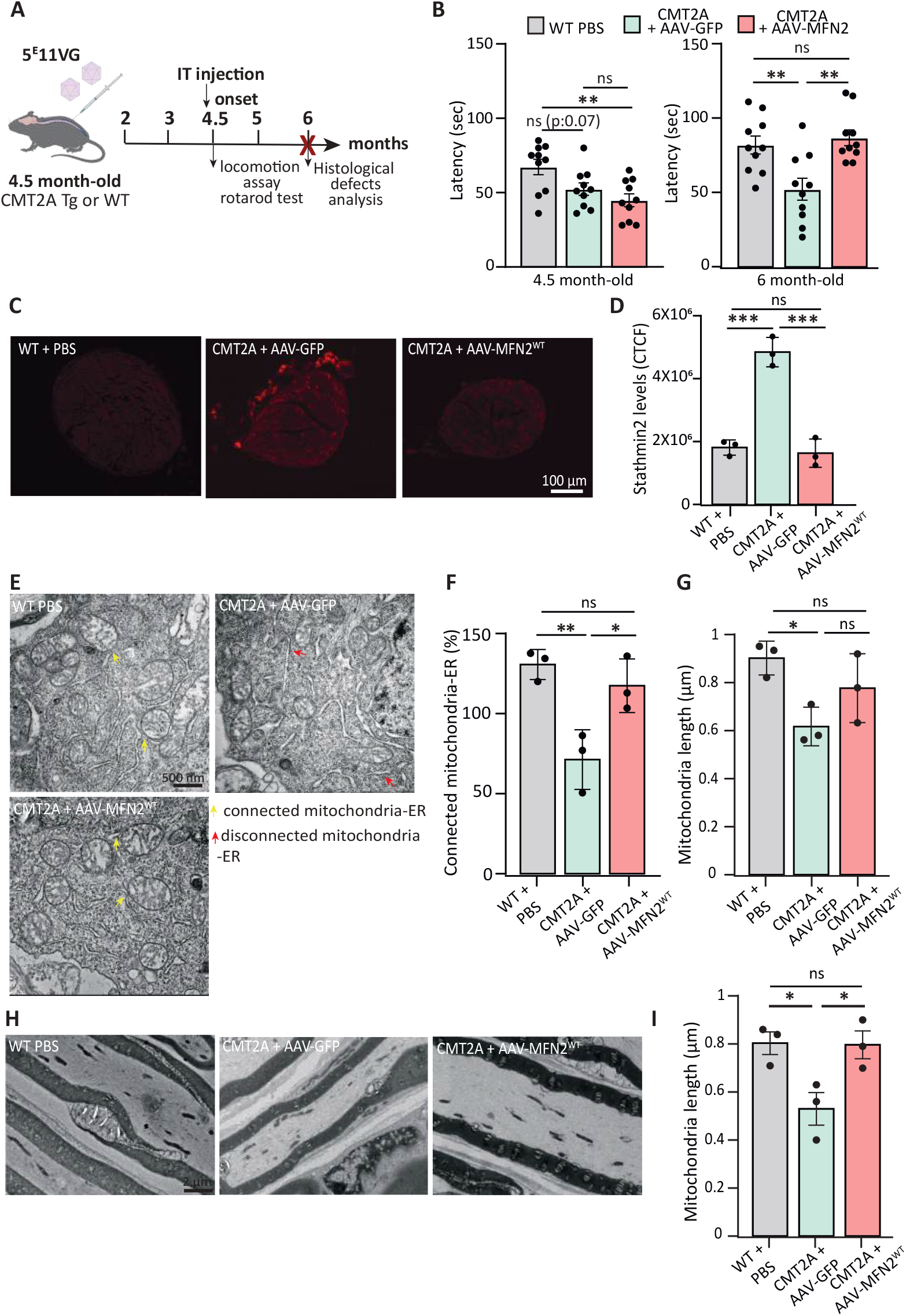
Post-symptomatic overexpression of MFN2 ^WT^ prevents locomotion defects and axonal degeneration. (A) Experimental design: at the onset (i.e. 4.5 months old), CMT2A transgenic (Tg) or wild-type (WT) mice received intrathecal injections of 5^E^11 vg AAV-hsyn-GFP or AAV-hsyn-MFN2^WT^. As control, WT mice were injected with PBS. Behavioral assessments and histological analyses were performed before the injection (4.5 months) and 6 weeks post-treatment (6 months). (B) Evaluation of rotarod performance measured at 4.5 and 6 months (n=10 mice for each of the experimental groups). Data are shown as mean ± SEM. Statistical analysis: one-way ANOVA multiple comparisons with Tukey’s post hoc test. (C) Representative immunofluorescence images of tibial nerve cross sections stained with anti-stathmin2 (red). (D) Quantification of stathmin2 expression levels using corrected total cell fluorescence (CTCF, n=3 animals per group). (E) Transmission electron microscopy (TEM) images of spinal cord motor neurons in the 3 conditions. Yellow arrowheads indicate mitochondria in close apposition to the endoplasmic reticulum (ER), whereas red arrowheads highlight disrupted mitochondria–ER contacts. (G) Quantification of mitochondria–ER apposition, expressed as the percentage of mitochondria closely associated with ER membranes relative to total mitochondria (H) Quantification of mitochondrial length on longitudinal sections of tibial nerves. For G and H, data are presented as mean ± SEM (n=3 animals per group). Statistical analyses were performed using one-way ANOVA multiple comparisons with Tukey’s post hoc test; *p* < 0.05 was considered significant.

### Intrathecal delivery of the therapeutic vectors does not cause liver or DRG toxicity

Previous gene therapy studies have highlighted the risk of hepatotoxicity of AAV vectors in mice, primates and humans^27,28^. In addition, intrathecal injection has been associated with transient DRG toxicity with certain promoters and at high doses^29^. Therefore, to determine if there were any toxic effects of our gene therapy approach, we analyzed several parameters indicating the aforementioned toxicity risks. The number of vector copies was quantified in various tissues (spinal cord, tibial and sciatic nerves, DRG and liver), revealing similar biodistribution in the mice injected with either AAV9-hsyn1-GFP or AAV9-hsyn1-MFN2^WT^. As expected, AAV9 showed high liver transduction for both vectors. The number of copies was higher in the spinal cord than in the DRG and the nerves (**Figure 5A**). This transduction profile after intrathecal (IT) injection confirmed the previously observed profile of transgene expression (**Figure 1)**. Importantly, despite high vector presence in the liver, there was not major visible signs of hepatotoxicity, such as cell death or cell infiltration or inflammation^27^. However, we noticed the presence of vacuolation mostly in the AAV9-hsyn1-GFP condition **(Figure 5B)**. We next assessed in DRGs the expression levels of ATF3, a marker of neuronal stress or injury, and IBA-1, a widely used marker of macrophages/microglia indicating immune cell infiltration. They both have been previously used as indicators of DRG injury following AAV gene therapies^30–32^. No significant changes in the number of ATF3^+^ or IBA-1^+^ cells were detected in DRGs up to 10 months after IT delivery of AAV9-hsyn1-GFP or AAV9-hsyn1-MFN2^WT^ compared to controls conditions injected with PBS (**Figure 5C-D and E-F)**.

**Figure 5.**
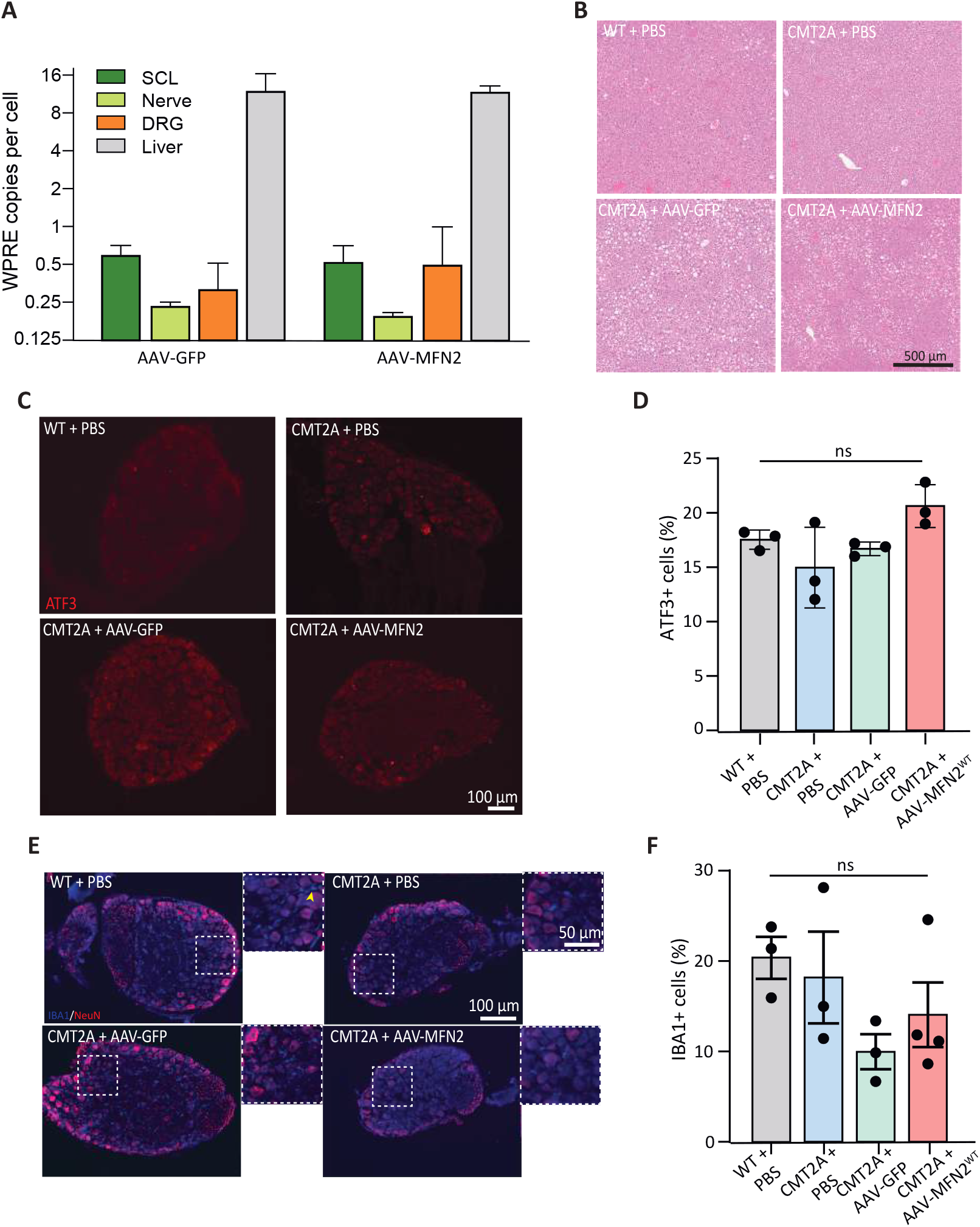
Intrathecal delivery of the therapeutic vectors does not lead to long-term hepatic or DRG toxicity. (A) Quantification of vector genome copies per cell (WPRE) in lumbar spinal cord (SCL), peripheral nerves (sciatic and tibial nerves combined), dorsal root ganglia (DRG), and liver, 10 months following AAV-GFP or AAV-hMFN2 administration. Data are presented as mean ± SEM (n=3 animals per condition). (B) Representative hematoxylin and eosin (H&E) staining of liver sections from WT and CMT2A mice treated with PBS, AAV-GFP, or AAV-hMFN2, showing no evidence of histopathological abnormalities. (C) Representative immunofluorescence images of DRG sections stained with anti-ATF3 (red) to assess neuronal stress in DRG in the different treatment conditions. (D) Quantification of the percentage of ATF3^+^ cells relative to NeuN^+^ cells in DRG do not show any statistical differences between the groups. (E) Representative immunofluorescence images of DRG sections stained with anti-IBA1 (blue) to label activated microglia/macrophages and counterstained with NeuN (red). Arrowhead indicate highlight IBA1+ cells. (F) Quantification of the percentage of IBA1+ cells relative to dapi^+^ cells in DRG does not show any change in microglial/macrophage activation after AAV administration compared with controls. For both D and F, data are presented as mean ± SEM (n=3 animals per condition). Statistical analyses were performed using one-way ANOVA multiple comparisons with Tukey’s post hoc test; *p* < 0.05 was considered significant.

## Discussion

Our study established the first in *vivo* proof of concept for gene therapy in CMT2A. In particular, we demonstrate that the overexpression of MFN2^WT^ in affected neurons is sufficient to rescue CMT2A pathology both before and after symptom onset. The current challenge in CMT2A lies in developing a therapeutic strategy that can target the vast majority of MFN2 mutations. More than 100 pathogenic variants have been reported in the *MFN2* gene, most of them being heterozygous autosomal dominant and only a minority being autosomal recessive^5–7^. The pathological effects of these mutations remain incompletely understood, as only a subset of them has been studied. A dominant-negative mechanism has been proposed, primarily for the most frequent and most studied mutation (MFN2^Arg94Gln^). In this case, the mutant form interferes with the WT protein, altering mitochondrial dynamics both *in vitro* (cell lines and hIPSC-derived MNs) and in rodent models^16,33^. Other *in vitro* studies suggest loss-of-function mechanisms, as the loss of MFN2 was sufficient to reproduce the deleterious effects of mutated MFN2 on mitochondrial dynamics or ER-mitochondria interaction^8,34^. In addition, the specific deletion of MFN2 *in vivo* in MNs led to muscle denervation, and muscle atrophy similar to those observed in CMT2A Tg mouse models^35^. On the contrary, recent work on Drosophila and cell lines further suggests the possibility of gain-of-function mechanisms for at least two MFN2 mutations (i.e., MFN2^Arg364Trp^ and ^Thr206Ile^), which, unlike MFN2^Arg94Gln^, promote mitochondrial fusion and ER-mitochondria contacts^12,13,36^. A recent cellular analysis of 12 MFN2 gene variants showed that only 6 of them significantly impaired mitochondrial fusion capacity. Most of these variants were located near or within the GTPase domain, and were associated with an early and severe form of the disease ^37^. Finally, structural studies have further revealed that MFN2 mutations alter the conformational dynamics of MFN2 and/or the formation of MFN2 homodimers, or MFN1/MFN2 heterodimers, in various ways, ultimately affecting GTP hydrolysis to a greater or lesser extent^38^. These differences may explain the various effects of the mutations observed in cellular and animal models.

Here, our strategy is based on gene supplementation (overexpression of MFN2^WT^), to increase the ratio of WT versus mutant protein levels. We believe this approach is sufficient to compensate for dominant-negative and loss of function effects which account for the majority of *MFN2* mutations. We indeed shown that this strategy is sufficient to counteract axonal degeneration in both hIPSC-derived MNs and CMT2A Tg mice (mitocharc1) as well as the associated locomotor deficits in these animals. Similar gene supplementation strategies have also been successfully applied to dominant-negative disease mechanisms such as inherited retinal dystrophies^39,40^. Furthermore, gene replacement/supplementation therapy currently represents the majority of clinically validated approaches for treating genetic diseases, accounting for 10 out of the 11 gene therapies approved by the US Food and Drug Administration^41^. An alternative strategy proposed for diseases associated with dominant-negative mutations, is to combine a knock-down of the causal gene followed by replacement with the WT version of the affected protein. This strategy was recently implemented in the context of CMT2A based on the use of RNA interference against MFN2 associated with the expression of exogenous MFN2 encoded by two separate AAV vectors^25^. The approach was successfully validated *in vitro* using hIPSC-derived MNs from CMT2A patients harboring the rare MFN2^Ala383Val^ mutation. However, the therapeutic benefit of this strategy was not evaluated *in vivo* on CMT2A mice.

Since *MFN2* mutations are mainly distributed across 3 hotspot regions (i.e. amino acid positions 94-105; 361-376; 705-740) (https://neuropathybrowser.zuchnerlab.net/) that contains the majority of *MFN2* mutations, prime-editing (PE) may be a relevant option to simultaneously target multiple CMT2A mutant alleles. Unlike the CRISPR–Cas9 approach, base editors and PE enable correction of pathogenic variants without generating double-strand breaks^42,43^. In particular, twin PE offers deletion and replacement of large DNA fragments (>100bp) and represents an innovative approach for genetic diseases associated with various mutations. However, most of the PoC studies using this technology has been conducted *in vitro* since delivery of the necessary components remains the current limitation for *in vivo* relying on the use of dual AAVs^44,45^. In addition, PE generally exhibits lower editing efficacy compared to base editing^46^.

Most of the current therapeutic development for CMT2A are focused on restoring mitochondrial fusion and distribution along the axons^20^. One of these approaches involved the development of synthetic compounds, agonists or activators, that promote a fusion-permissive conformation of MFN2 by destabilizing the HR1 and HR2 domains of MFN2^15,47^. These compounds showed positive effects by reversing defective mitochondrial dynamics in hIPSC-MNs carrying the p.Arg94Gln or p.

Thr105Met mutations in MFN2 and in CMT2A mice expressing MFN2^Thr105Met^. However, the need of repeated administrations and the risks associated with constitutive mitochondrial fusion activation remain a major challenge for clinical translation. Another strategy is based on the overexpression of MFN1, the homolog of MFN2^15,16^. By crossing Tg Prp-MFN1 mice with Thy1.2-MFN2^Arg94Gln^ CMT2A mice, Zhou and coll. demonstrated that overexpression of MFN1 prevents the development of CMT2A pathology. Whether MFN1 overexpression could be protective when induced at later stages has not yet been assessed^16^. MFN1 supplementation has also been tested *in vitro*, either in primary sensory neurons expressing MFN2^Arg94Gln^ or more recently in SH-SY5Y cells expressing the MFN2^Lys357Thr^ mutation. The authors showed that MFN1^WT^ is more effective than MFN2^WT^ in preventing the mitochondrial clustering observed in CMT2A-SH-SY5Y cells^18^. Only a partial rescue of mitochondrial dynamics has been observed in CMT2A-sensory neurons^48^. Surprisingly, in our initial study published in 2019, we did not observe any rescue on axonal defects by overexpressing MFN1 in primary CMT2A mouse MNs^11^. The finding that MFN1, which is normally restricted to the mitochondrial membrane and regulates mitochondrial fusion, failed to reproduce the therapeutic effect of MFN2^WT^ led us to hypothesize that MFN2’s beneficial role may arise from its additional functions, particularly the regulation of ER-mitochondria contacts. Since our initial study, other groups have reported defective contacts between these organelles in different cellular models and with different MFN2 mutations^12–14^. These contacts, also known as mitochondria-associated membranes (MAMs), regulate essential neuronal functions, including lipid and calcium exchange, autophagy/mitophagy, and mitochondrial dynamics^11,49^, for which long-projecting neurons, such as motor and sensory neurons, are particularly dependent. Interestingly, a recent study identified MFN2 transcripts variants encoding shorter isoforms of the protein^9^. These isoforms are either localized at the ER membrane (ERMIN2) or act as a link between the ER and mitochondrial membranes (ERMIT2). When overexpressed, ERMIT2 promotes ER–mitochondria juxtaposition by interacting with MFN1 or 2 via its coiled-coil domain. Given that CMT2A is associated with a predominantly motor phenotype^5^, we aimed to develop a tool capable of efficiently targeting lumbar spinal MNs, which innervate the distal limbs, the most severely affected regions in both patients and our mouse model. We selected AAV9 because it is currently the most commonly vector used to treat neurological disorders thanks to its ability to cross the blood–brain barrier and efficiently transduce neuronal tissue. Notably, it was also the first AAV serotype to be approved as a medical product for the treatment of spinal muscular atrophy (Zolgensma®). We opted for IT over IV delivery because it targets the lumbar spinal cord leading to high MNs transduction. In addition, IT delivery may reduce side effects by bypassing the systemic circulation and the risk of inhibition by circulating neutralizing antibodies^50^. Another advantage of IT injection is that it is minimally invasive compared with procedures such as subpial injection^51^ and it is already used in clinics. In the field of axonal CMT, some gene replacement therapies have been developed and are currently in clinical trials. Notably, these include IGHMBP2 gene replacement for SMARD1 (MIM 604320) and CMT2S (MIM 616155) delivered via IV or ICV injection^52,53^ (NCT05152823) as well as restoration of gigaxonin expression in giant axonal neuropathy (GAN, MIM 256850) using IT administration of scAAV9 (NCT02362438)^24,54^. For this clinical trial, the first results were encouraging, showing an improvement of motor score one year after injection. Unfortunately, among the 14 participants, 2 individuals who received the lowest dose died unexpectedly due to post-operative complications. Although gene therapy is increasingly used to treat incurable and rare diseases^55^, adverse effects and potential toxicity remain a concern in AAV-based approaches, particularly when local targeting cannot be achieved. These issues underscore the need for careful risk–benefit assessment in the development of AAV-based gene therapeutics^56,57^, and highlight the importance of ongoing efforts to engineer safer vectors.

One promising approach to improve safety involves the use of cell-type specific promoters or enhancers which can restrict transgene expression to the affected population and reduce the risk of off-target effects. This strategy has already been successfully implemented in demyelinating form of CMT using the Mpz/P0 promoter^22,58^. In our study, despite local intrathecal injection IT AAV9 injection resulted in high levels of AAV genome copies in the liver. It would be important to include future comprehensive liver functions studies to complement the histopathological findings. Furthermore, concerns have recently been raised regarding DRG toxicity caused by IV or IT administration of AAV in non-human primates (NHPs)^59^, and in patients with amyotrophic lateral sclerosis (ALS) and GAN ^60,61^, potentially leading to chronic pain. In our model, we observed no evidence of DRG toxicity following AAV9 administration, as assessed by IBA-1 (an indicator of microglial/macrophage invasion) and ATF3 (a marker of neuronal injury)^30–32^ stainings. One possible solution to further limit the expression of the transgenes in DRGs could be to include of microRNA-target sequences ^62^. In parallel, recent optimizations of AAV capsids and cis regulatory elements have been shown to improve MN targeting with limited liver or DRG targeting after IV or IT injections in rodent and non-human primate models^63–65^. These innovations open new and potentially safer therapeutic avenues in the field of MN diseases.

Finally, an important limitation to address is the large number of CMT2A subtypes arising from different MFN2 mutations. Only a few rodent models of CMT2A have been developed, outside the one with the p.Arg94Gln mutation and very few display a robust CMT2 phenotype, limiting our ability to assess the efficacy of our therapy on other mutations. Recently, two rat knock-in models harboring MFN2^Arg364Trp^ or MFN2^His361Tyr^ have been generated (https://www.psychogenics.com/resources/characterization-of-a-rat-model-of-cmt2a/ and ^26^), both exhibiting a predominant motor phenotype. These models will be useful for the testing of future therapies. At the same time, hIPSC-derived motor neurons and neuromuscular organoids carrying various MFN2 mutations^66,67^ represent valuable models for testing therapies in a mutation-specific context. Combined with *in vivo* studies, they will be critical to assess the generalization of our gene supplementation strategies and for designing treatments that address the mechanistic variability of CMT2A. In conclusion, we have established the first *in vivo* proof of concept for gene therapy applicable to CMT2A, demonstrating that MFN2^WT^ overexpression restores axonal integrity by rescuing ER–mitochondria contacts. These preclinical results will pave the way for future AAV-based treatments for CMT2A in clinics as well as for other neuropathies and motor neuron diseases linked to mitochondrial-ER dysfunction.

## Acknowledgements

We acknowledge the PICsL core facilty (IBDM, AMU-Marseille), in particular Fabrice Richard, who performed the electron microscopy experiments. We are grateful to the PTBTG platform (EPFL, Lausanne, Switzerland), in particular Aline Aebi, Jean-Philippe Gaudry and Philippe Colin for vector production, genome copy number quantification and intrathecal injection training.

## Fundings

This research work was supported by the French National Research Agency (ANR) JCJC grant: #ANR-21-CE17-0033-01 and the starting grant INSERM).

## Competing Interests

The Authors declare that there is no conflict of interest.

## Supplementary Material

Supplementary Figure is provided as separate PDF files.

## Abbreviations

AAV: Adeno-associated virus
CMT: Charcot-Marie-Tooth
ER: endoplasmic reticulum
hsyn1: human synapsin
IT: intrathecal
IV: intravenously
MFN2: mitofusin-2
MN: motor neuron
MAM: mitochondria-associated membrane
PE: prime-editing.

## Notes

### Competing Interest Statement

The authors have declared no competing interest.

### Summary of Updates

The margins have been adjusted in all figures.

