## supplementary figure1 for "Gene therapy-mediated overexpression of wild-type MFN2 improves Charcot-Marie-Tooth disease type 2A"

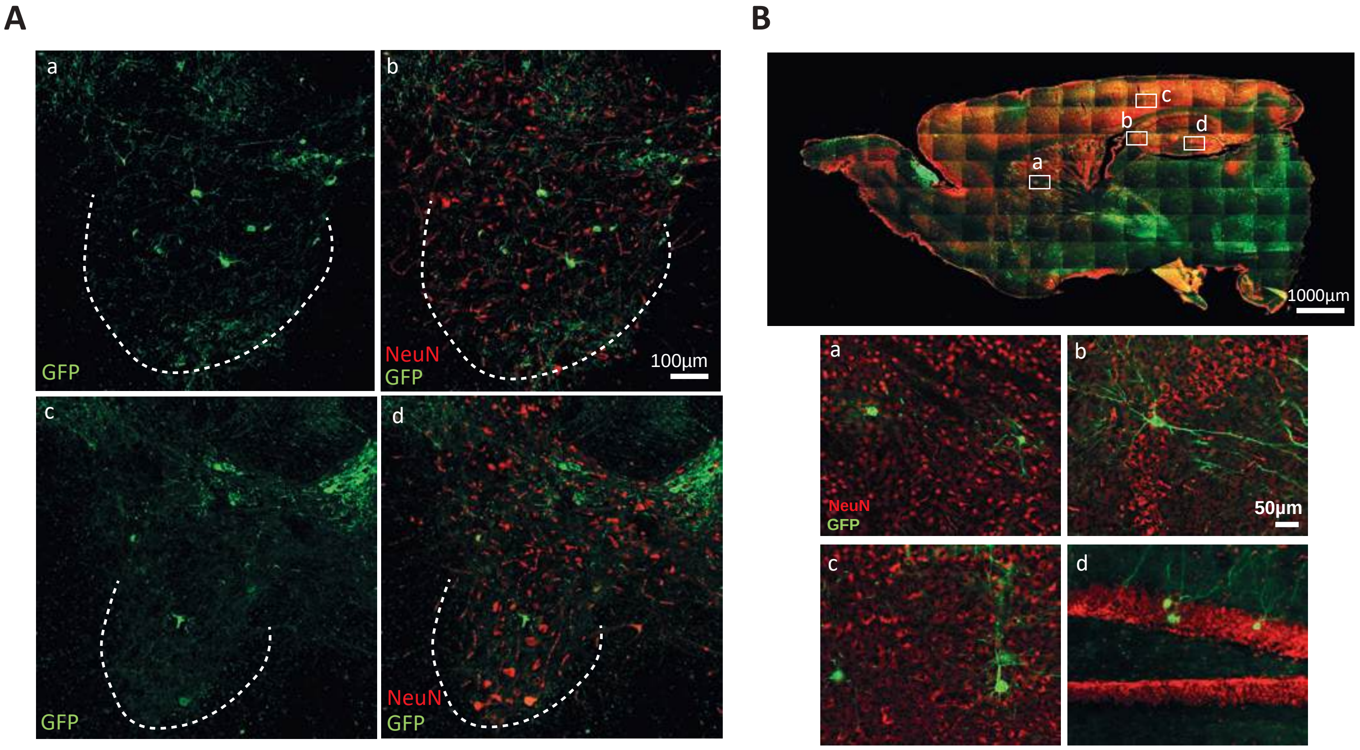
A
a
b
GFP
NeuN
GFP
100µm
c
d
GFP
NeuN
GFP
B
c
b
d
a
1000µm
a
b
NeuN
GFP
50µm
c
d

**Figure.S1**

**FigureS1. Biodistribution and CMT2A line characterization**

(A) Coronal sections of the spinal cord showing cell transduction in cervical (upper panels, a-b) and thoracic (lower panels, c-d) regions, showing GFP expression (green), and GFP/NeuN merged staining. Scale bars: 100 µm. (B) Sagittal brain sections illustrating GFP biodistribution (green) and NeuN labeling (red). Higher magnifications of selected regions highlight neuronal transduction in (a) lateral septal nucleus, (b) hippocampal CA2 region, (c) cerebral cortex, and (d) dentate gyrus. Scale bars: 1000 µm (overview), 50 µm (insets).
